# Short-term passive greenspace exposures have little effect on nasal microbiomes: a cross-over exposure study of a Māori cohort

**DOI:** 10.1101/2024.01.17.576148

**Authors:** Joel E. Brame, Isaac Warbrick, Deborah Heke, Craig Liddicoat, Martin F. Breed

## Abstract

Indigenous health interventions have emerged in New Zealand aimed at increasing human interactions with and exposure to macro and microbial diversity. Urban greenspaces provide opportunities for people to gain such exposures. However, the dynamics and pathways of microbial transfer from natural environments onto a person remain poorly understood. Here, we analysed bacterial 16S rRNA amplicons in air samples (*n* = 7) and pre- and post-exposure nasal samples (*n* = 238) from 35 participants who had 30-minute exposures in an outdoor park. The participants were organised into two groups: over eight days each group had two outdoor park exposures and two indoor office exposures, with a cross-over study design and washout days between exposure days. We investigated the effects of participant group, location (outdoor park vs. indoor office), and exposures (pre vs. post) on the nasal bacterial community composition and three key suspected health-associated bacterial indicators (alpha diversity, generic diversity of Gammaproteobacteria, and read abundances of butyrate-producing bacteria). The participants had distinct nasal bacterial communities, but these communities did not display notable shifts in composition following exposures. The community composition and key health bacterial indicators were stable throughout the trial period, with no clear or consistent effects of group, location, or exposure. We conclude that 30-minute exposure periods to urban greenspaces are unlikely to create notable changes in the nasal microbiome of visitors, which contrasts with previous research. Our results suggest that longer exposures or activities that involves closer interaction with microbial rich ecological components (e.g., soil) are required for greenspace exposures to result in noteworthy changes in the nasal microbiome.

## INTRODUCTION

Disconnection from natural environments is a characteristic of urban lifestyles and one which is associated with poorer health outcomes (1, 2). For Indigenous Peoples, whose identity, culture, and health are intertwined with the natural environment (3, 4), the disconnection from ancestral lands and natural environments generally, is particularly concerning. Warbrick et al (2023) recently proposed that the relationship between environmental microbiomes and health has important implications for the health of Indigenous Peoples, despite Indigenous people rarely being represented in studies of the microbiome. With the majority of people now living in cities (5), urban greenspaces and their accompanying aerobiomes are key points of exposure to natural environmental microbiomes (6).

Bacterial colonisation of the human body occurs during and after birth, with post-birth bacterial communities primarily shaped by people’s environments (7). Pathways of exposure to environmental bacteria include ingested and inhaled substances, either directly or indirectly (e.g., via hand-to-face transfer). Air is a well-understood transmission medium for microbiota, which triggers health conditions such as allergies and infectious disease (8). However, the transmission pathway of health-supporting airborne bacteria has received much less attention (6). Airborne bacterial communities (aerobiomes) of built indoor environments are highly variable due to a wide range of possible conditions (9). Outdoor environments are also rich aerobiome reservoirs (10). Because airborne dispersal of microbiota is a key pathway of bacterial exposure and transfer, air transfer dynamics can be studied via sampling nasal bacterial communities (6). Nasal microbiome changes may reflect the characteristics of aerobiomes of recent exposure, suggesting that the study of outdoor aerobiomes can provide critical insights into human microbiome assemblages (11). However, few studies have examined how nasal microbiomes change after exposure to outdoor air.

Greenspace aerobiomes originate from leaf surfaces and soil, with modulating effects from vegetation complexity and height above the ground (i.e., vertical stratification; 10), air pollution (12), and wind-carried airshed influences (6). In urban settings, land cover has a strong influence on the composition of aerobiomes. For example, the aerobiomes of parks have different community compositions than adjoining parking lots (13). Among greenspaces, amenity grassland aerobiomes have different compositions to remnant native vegetation aerobiomes and possess consistent alpha diversity at heights up to 2 m (10). Thus, urban amenity grasslands should have distinct aerobiomes compared to indoor offices and provide useful locations to study the transfer of aerobiomes into the airways of people. Yet, the use of amenity grassland aerobiomes in bacterial transfer studies is limited.

Several aerobiome characteristics and taxonomic groups have been linked with human health. Salutogenic functions of bacteria include maintenance of the mucosal barriers (14), immune signalling (15), vitamin production (16), and synthesis of short-chain fatty acids such as butyrate (17). The Biodiversity Hypothesis describes how exposure to a greater amount of microbial diversity in the natural environment may be required to promote innate immune training and immunoregulation (18). In a complex network of interactions, exposure to bacterial diversity can modulate immune responses and reduce pro-inflammatory and allergenic antibodies and cytokines (18). Thus, exposure to higher alpha diversity of bacteria within outdoor aerobiomes with a low level of pathogenic taxa could potentially support human health (19).

The diversity of Gammaproteobacterial genera on the skin has been associated with increased plasma transforming growth factor beta 1 (TGF-β1) levels, decreased interleukin-17 (pro-inflammatory cytokines), and increased relative abundance of regulatory T-cells (20). Increased TGF-β1 and decreased interleukin-17 are associated with an anti-inflammatory molecular profile, and regulatory T-cells are critical for immunotolerance, including tolerance of commensal taxa (20). Butyrate-producing bacteria are key members of the human and animal gut with numerous health benefits, and after birth they are primarily supplied by the environment with nutritional support via ingestion of fibre (17). Certain outdoor environments are reservoirs of butyrate producers that could disperse into the aerobiome and transfer onto people visiting those environments (21). Thus, aerobiome Gammaproteobacterial diversity and butyrate-producing bacterial read abundances could provide indicators of human health-associated benefits of aerobiome exposure.

Here we studied the changes in 16S rRNA amplicons in pre- and post-exposure nasal microbiome samples from 35 Māori (Indigenous New Zealand) participants, divided into groups (A and B), who spent two repeated 30-minute exposure periods in each of two locations: an indoor office and an outdoor park (amenity grassland). We utilised a cross-over study design to control for effects of group and day, with two exposure days in one location (Days 1 and 3), followed by a two-day washout period, then two further exposure days in the other location (Days 6 and 8). To understand the influences of exposures on the nasal microbiomes, we examined the effects of location, individual, group, single exposures, and repeated exposures on (1) the nasal bacterial alpha diversity, (2) nasal bacterial community composition, and (3) specific bacterial taxonomic groups with known health associations (Gammproteobacterial diversity and butyrate-producing bacterial abundances).

## MATERIALS AND METHODS

### Experimental design

We utilised a crossover trial design (**Figure 1A**). We recruited 35 participants into the trial, which took place March 15-22, 2023. The participants were divided into two groups: outdoor and indoor for exposure days 1 and 3, with crossover for exposure days 6 and 8. Exposure days were on March 15 and 17, then on March 20 and 22, allowing for a single washout day between testing days and two washout days before the crossover.

**Figure 1.**
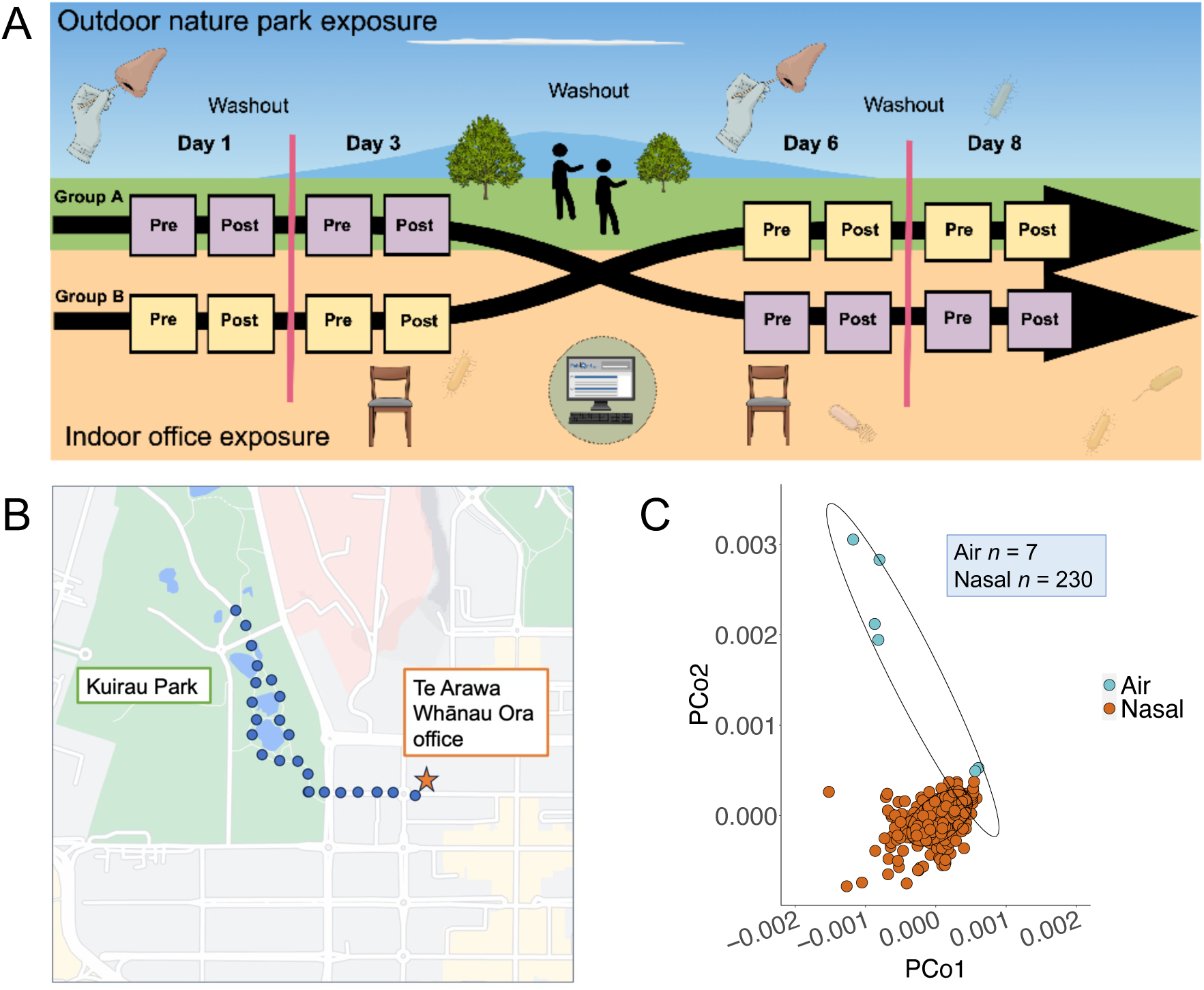
(A) Overview of the cross-over experimental design. (B) Walking path map of the outdoor treatment group in Rotorua, New Zealand. Map generated with Google maps. (C) Principal coordinates analysis based on centred-log ratio compositional abundance data displaying variation in community composition by sample type (Adonis PERMANOVA: F = 5.515, R^2^ = 0.023, *p* = 0.001).

The outdoor treatment group met at the Te Arawa Whānau Ora office in Rotorua, New Zealand, at approximately 8:30am. Te Arawa Whānau Ora is an Indigenous community health organisation, and all participants in this study were employees of the organisation and identify as Māori. Their noses were swabbed pre-exposure (hereafter referred to as “Pre”, see description below), and they then went for a walk to Kuirau Park, approximately 600 m from the office, for 30 minutes (**Figure 1B**). Upon their return, before entering the office, they were re-tested with a second nasal swab (hereafter referred to as “Post”).

The indoor treatment group met at the same office at the same time and day as the outdoor treatment group. Their noses were swabbed using the same methods. However, during the 30-minute exposure period, they remained in the office.

### Nasal swabbing

Nasal swab samples were obtained by inserting a sterile nylon-flocked swab tip (FLOQSwabs Lot 2011490, Copan Flock Technologies, Bescia, Italy) into the anterior nares and rotating in a circular motion for 3-5 seconds per naris, then repeated in the opposite naris using the same swab. The swab tip was then immediately snipped into a sterile 15 mL falcon tube, sealed with the lid, wrapped with parafilm, and placed in a -20°C freezer in the office.

### Air sampling

Air samples were obtained at an outdoor park site along the same walking path where participants walked during their outdoor period and at a central indoor location in the Te Arawa Whānau Ora office. The Kuirau Park site in Rotorua, New Zealand is predominantly amenity grassland with interspersed geothermal springs. At Kuirau Park, air samples were collected over an approximately 8-hour period during each testing day, following the method described in Mhuireach, Johnson (22). The aerobiome sampling stations were set up on site between 0800 and 0830 hours and collected between 1500 and 1530 hours. At the Te Arawa Whanau Ora office control site, air samples were collected following the same procedures and the same times. On one day, March 17, the weather was rainy and the air stand assembly using protective umbrellas was vandalised, thus an outdoor air sample was not obtained for that day.

The outdoor park air sampling station was made of plastic boxes and achieved a height of 1.2 m. Sampling at this height should be representative of aerobiome exposure potential for children and adults alike, and is within the 2 m height range of similar alpha diversity as measured elsewhere in amenity grassland aerobiomes (10). The indoor office sampling station was a single plastic box placed on a table, achieving a height of approximately 1.5 m. On the top of each station, we opened and placed three sterile clear plastic petri dish bases and lids, which provided six collection surfaces per site. This method of passive aerobiome sampling has been shown to be as effective as active sampling methods (22). On two days, a field control was generated by holding open an additional petri dish for 30 seconds at the equipment box. Immediately after the air sampling activity, each petri dish was sealed, labelled, and placed in the office freezer at -20°C until DNA extraction (described below).

### DNA extraction, PCR, sequencing

Within one week of obtaining all samples, DNA extractions and quantifications were performed in a PC2 laboratory at Auckland University of Technology, Auckland, New Zealand. To transport samples from Rotorua to Auckland, samples were removed from the office freezer, placed onto ice in a sealed insulated container, and transported by vehicle to the lab. Upon arrival at the lab, they were immediately placed into a -20 °C freezer.

The petri dishes for each site were opened and swabbed with sterile nylon-flocked swab tips (FLOQSwabs) inside a laminar flow cabinet. One swab and 40 uL of added sterile phosphate-buffered saline was used for swabbing all six surfaces, except for surfaces that showed visual signs of damage or contamination, for approximately four minutes total using a consistent pattern of swabbing. The tips were cut directly into 15 mL sterile falcon tubes. We obtained an extraction blank control for each extraction batch using the same process as samples but without a swab tip.

We used the QIAamp DNA Mini Kit (QIAGEN) for all samples and followed the manufacturer’s instructions with two modifications to increase final concentration: the incubation step was extended from 10 min to 15 min, and the final elution buffer volume was reduced from 80 uL to 60 uL. The extraction concentrations were then quantified using a Qubit High Sensitivity dsDNA assay (ThermoFisher Scientific). Once DNA concentrations were verified, PCR amplification of the bacterial 16S rRNA V3-V4 regions was performed in the lab at Auckland University of Technology using Kappa HiFi Taq mix with 341F-805R primers (Kapa Biosystems) via PCR on an Eppendorf Vapo.Protect Mastercycler Pro thermocycler. The first PCR round included 38 amplification cycles. Plate clean-up was performed via AMPure XP reagent. To normalise clean PCR products to 1 ng/uL, samples below 1 ng/µL were concentrated using the Eppendorf Concentration and using the following conditions: D-AQ, 30 C, 18 min. Second round PCR used the Nextera XT Index Kit to index samples, with eight cycles of amplification. Samples were then pooled, cleaned with AMPure XP reagent, and quantified using Qubit High Sensitivity. The Bioanalyzer 2100 expert High Sensitivity DNA assay was performed to check library quality and molarity, and libraries were pooled for equal molarity. Upon completion of library preparation, sequencing of amplicon sequence variants was completed on the Illumina Miseq V3 using the Illumina MiSeq Reagent Kit v3 (600 cycle). Four PCR negative blanks were generated during the library preparation steps for quality control.

### Bioinformatics

From the 16S rRNA raw sequence data, amplicon sequence variants (ASVs) were trimmed and filtered using an established Qiime2 pipeline (version 2023.5), with forward reads truncated at 260 bp and reverse reads truncated at 198 bp. Taxonomy was assigned using the onboard Naïve Bayes taxonomic classifier and Silva database v 138.1. Sequences were then cleaned using scripts utilising the R *phyloseq* package (version 1.42.0; 23) by removing the following sequences: those assigned to mitochondria and chloroplasts, taxa that did not occur in at least two samples, and ASVs with total sums < 20 reads. Sequences that were likely of contamination origin were identified and removed using the R *decontam* package (version 1.18.0; 24) using the function “isNotContaminant” suited for low biomass samples.

### Statistical analysis

All statistical analyses were performed using R (version 4.2.3; 25). To maintain consistency with prior aerobiome studies, statistical significance was set at alpha = 0.05. Sample alpha diversity based on Hill numbers was examined using R *hillR* package (26), which integrates sample size and coverage. We set the q parameter for Hill numbers at 0.80 for reduced sensitivity to relative abundances compared with Shannon index.

To prepare for beta diversity tests, the read abundance data were evaluated using R *zCompositions* package (version 1.4.0.1; 27), zeros were imputed using the R *scImpute* package (version 0.0.9; 28), and eight low total read abundance samples were discarded to reduce data sparsity. The resultant read abundances were then transformed with centred-log ratio using the R *compositions* package (verion 2.0.6; 29), followed by ordination with principal coordinates analysis using R *ecodist* package (version 2.0.9; 30), based on Aitchison distances obtained with the R *vegan* package (version 2.6.4; 31); statistics were generated using PERMANOVA (Adonis) tests via the R *vegan* package. Distance-to-centroid analyses were performed using the R *vegan* package. Maps were created using the R *ggmap* package (version 3.0.2; 32). Differential abundance analysis using the Analysis of Compositions of Microbiomes with Bias Correction (ANCOM-BC) method was performed on untransformed amplicon data with the *ancombc2* function in the R *ANCOMBC* package (version 2.0.3; 33). Participant was set as a random effect (rand_formula) for mixed effects modelling. The p-value adjustment was set as “fdr”, and prv_cut and lib_cut were set at “0”. The ANCOMBC algorithm has been shown to minimise bias due to sampling fractions and reduces false discovery rates. We downloaded a comprehensive list of pathogens from Bartlett, Padfield (34)) to examine pathogenic read abundances in the samples. Time-series analyses were performed using repeated-ANOVAS with R *rstatix* package (version 0.7.2; 35). R *ggplot2* package (version 3.4.2; 36) was used for data visualisations.

## RESULTS

### Aerobiomes were different from nasal microbiomes

Aerobiomes had a higher alpha diversity (hill number = 82.3 ± 64.4 SD, *n* = 7) than nasal microbiomes (hill number = 19.5 ± 10.6 SD, *n* = 238; W = 1378, *p* = 0.003). Aerobiome location had no effect on alpha diversity between outdoor (hill number = 60.4 ± 70.3 SD, *n* = 3) and indoor (hill number = 98.8 ± 64.6 SD, *n* = 4) samples (t = 0.740, df = 4.2, *p* = 0.50). Overall, aerobiomes and nasal microbiomes had different community compositions (Adonis PERMANOVA: F = 5.515, R^2^ = 0.023, *p* = 0.001; **Figure 1C**), and outdoor aerobiomes were compositionally similar to indoor aerobiomes (Adonis PERMANOVA: F = 1.268, R^2^ = 0.20, *p* = 0.17).

### Exposure effect on composition, diversity and differential ASV abundances

The 30-minute outdoor exposures did not change the nasal bacterial community composition for either group A (Adonis PERMANOVA: F = 0.686 R^2^ = 0.013, *p* = 0.99) or group B (Adonis PERMANOVA: F = 0.726, R^2^ = 0.013, *p* = 0.98) (**Figure 2A**). The 30-minute indoor exposures also did not change the community composition for either group A (Adonis PERMANOVA: F = 0.809, R2 = 0.014, *p* = 0.89) or group B (Adonis PERMANOVA: F = 0.675, R^2^ = 0.012, *p* = 0.99) (**Figure 2D**).

**Figure 2.**
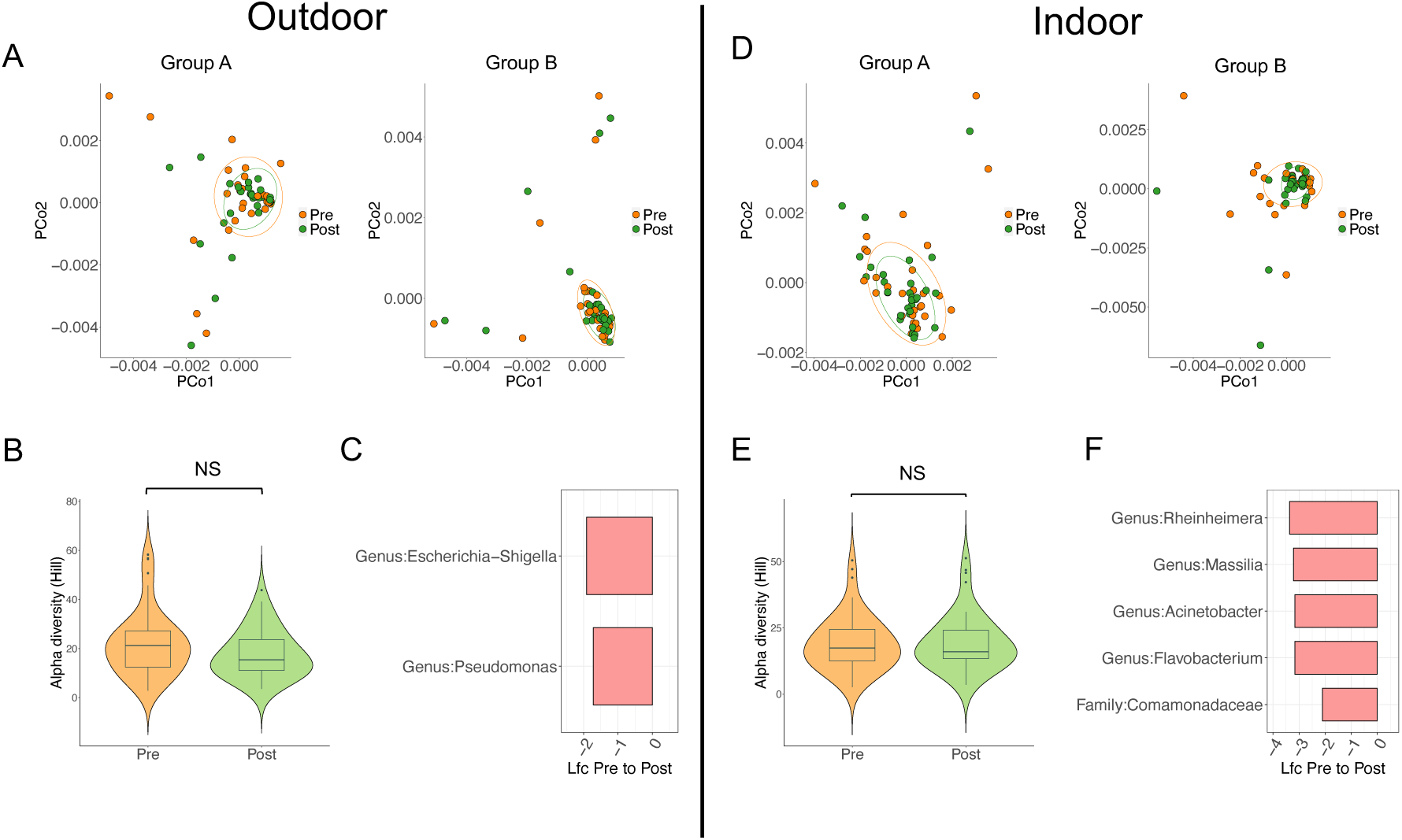
(A) Principal coordinates analysis based on centred-log ratio compositional abundance data displaying variation in community composition before (Pre) and after (Post) outdoor exposure for groups A and B. (B) Boxplots of changes in alpha diversity from before (Pre) and after (Post) outdoor exposure. The y-axis shows the alpha diversity based on Hill numbers. Boxes show the median and interquartile range, while whiskers extend to the remaining range of data. (C) Significantly differentially abundant genera in nasal microbiomes after outdoor exposure. The x axis shows the log fold change from before pre-exposure to post-exposure. Red bars indicate a decrease in log fold change. (D) Principal coordinates analysis based on centred-log ratio compositional abundance data displaying variation in community composition before (Pre) and after (Post) indoor exposure for groups A and B. (E) Boxplots of changes in alpha diversity from before (Pre) and after (Post) indoor exposure. The y-axis shows the alpha diversity based on Hill numbers. Boxes show the median and interquartile range, while whiskers extend to the remaining range of data. (F) Significantly differentially abundant genera in nasal microbiomes after indoor exposure. The x axis shows the log fold change from pre-exposure to post-exposure. Red bars indicate a decrease in log fold change.

There was no effect of group on changes in nasal bacterial alpha diversity after 30-minute exposures among both outdoor exposures (Wilcox: W = 433, p = 0.83) and indoor exposures (Wilcox: W = 358, p = 0.18), even though the groups visited the locations on separate days. When the two groups were combined, there was no effect on the alpha diversity by either the outdoor exposures (W = 1388, *p* = 0.11; **Figure 2B**) or the indoor exposures (W = 1818, *p* = 0.93; **Figure 2E**), although the treatment location effect (i.e., indoor vs outdoor) was significant (Wilcox: W = 2169, *p* = 0.02).

For group A, the 30-minute outdoor exposure had no effect on the read abundance of any genus on day 1. However, on day 3, the outdoor treatment resulted in a significant decrease in the genera *Escherichia-Shigella* (ANCOMBC: log fold change (lfc) = -1.91, adjusted-*p* (*q*) < 0.001) and *Pseudomonas* (ANCOMBC: lfc = -1.72, *q* < 0.001) (**Figure 2C**). For group B, on day 6, the outdoor treatment resulted in five taxa with significantly decreased read abundances: *Rheinheimera* (ANCOMBC: lfc = -3.37, *q* < 0.001), *Massilia* (ANCOMBC: lfc = -3.22, *q* < 0.001), *Acinetobacter* (ANCOMBC: lfc = -3.16, *q* < 0.001), *Flavobacterium* (ANCOMBC: lfc = -3.16, *q* < 0.001), and family Comomonadaceae (ANCOMBC: lfc = - 2.10, *q* = 0.004; **Figure 2F**). The outdoor treatment had no effect on any genus on day 8 for group B (all data are in **Table S1**).

### Exposure effects on health-associated bacterial groups

30-minute exposures had different effects in groups A and B on the number of Gammaproteobacteria genera (t-test: t = -2.111, df = 115.12, *p* = 0.036), so we examined the two groups separately. Indoor exposure significantly decreased the Gammaproteobacteria diversity in group A (t-test: t = -2.221, df = 56.61, *p* = 0.03) but had no effect in group B (Wilcox: W = 358, *p* = 0.91). Outdoor exposure had no effect on Gammaproteobacteria diversity for group A (t-test: t = -1.015, df = 49.3, *p* = 0.32) but weakly decreased Gammaproteobacteria diversity for group B (t-test: t = -1.905, df = 54.99, *p* = 0.062).

There was no effect of group on changes in nasal butyrate-producing bacterial read abundances after 30-minute exposures among both outdoor exposures (Wilcox: W = 329, *p* = 0.24) and indoor exposures (Wilcox: W = 396.5, *p* = 0.43). With Groups A and B combined, we observed no effect of treatment location on butyrate producer read abundances (Wilcox: W = 1940, *p* = 0.28).

### Aerobiome-associated taxa in nasal microbiomes

We identified 1098 bacterial taxa in the outdoor aerobiome samples and then constrained nasal microbiome analyses with only these taxa. 30-minute exposures had no effect on the percentage of aerobiome taxa in nasal samples in either outdoor (t-test: t = 0.331, df = 114.77, *p* = 0.74; **Figure 3A**) or indoor treatments (W = 1740, *p* = 1; **Figure 3D**). 30-minute outdoor exposures had no effect on the community composition of aerobiome taxa in nasal samples in either group A (Adonis PERMANOVA: F = 0.761, R^2^ = 0.014, *p* = 0.96; **Figure 3B**) or group B (Adonis PERMANOVA: F = 0.861, R^2^ = 0.015, *p* = 0.77; **Figure 3C**), and 30-minute indoor exposures had no effect on the community composition of aerobiome taxa in nasal samples taxa in either group A (Adonis PERMANOVA: F = 0.862, R^2^ = 0.015, *p* = 0.80; **Figure 3E**) or group B (Adonis PERMANOVA: F = 0.704, R^2^ = 0.013, *p* = 0.99; **Figure 3F**).

**Figure 3.**
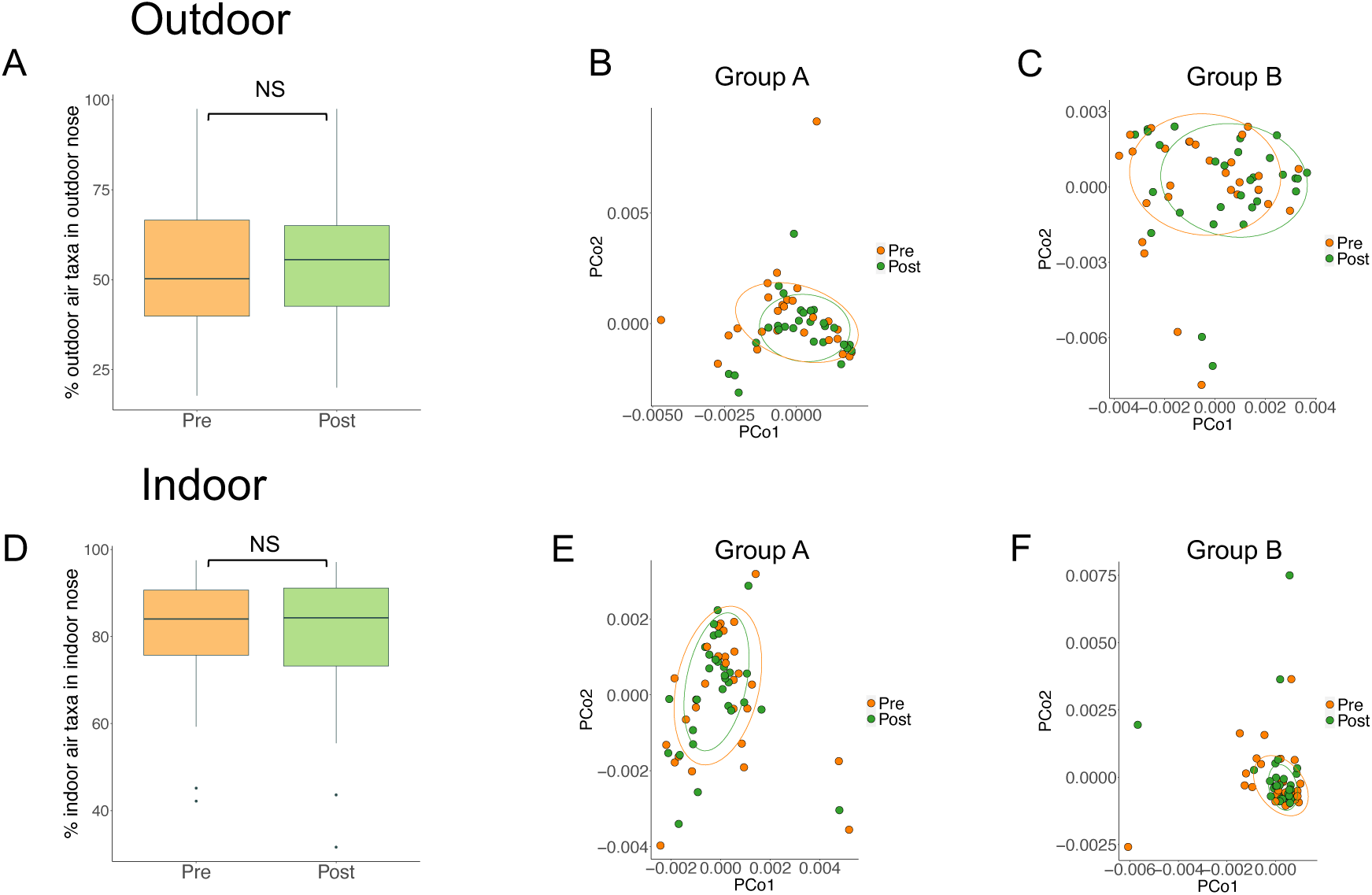
(A) Boxplots of the percentage of outdoor air taxa that were found in the nose (y-axis) before (Pre) and after (Post) outdoor exposure. Boxes show the median and interquartile range, while whiskers extend to the remaining range of data. (B-C) Principal coordinates analysis based on centred-log ratio compositional abundance data of only aerobiome-associated taxa found in the nose, displaying variation in community composition before (Pre) and after (Post) outdoor exposure for groups A (panel B) and B (panel C). (D) Boxplots of the percentage of outdoor air taxa that were found in the nose (y-axis) before (Pre) and after (Post) outdoor exposure. Boxes show the median and interquartile range, while whiskers extend to the remaining range of data. (E-F) Principal coordinates analysis based on centred-log ratio compositional abundance data of only aerobiome-associated taxa found in the nose, displaying variation in community composition before (Pre) and after (Post) outdoor exposure for groups A (panel E) and B (panel F).

### Time-series effects on nasal microbiome characteristics

Participant had a strong effect on nasal bacterial communities from Day 1 to Day 8 (Adonis PERMANOVA: F = 3.667, R^2^ = 0.382, *p* = 0.001; **Figure S1**). However, time had no effect on post-exposure group nasal bacterial community composition (**Figure 4A-D**). Group homogeneity (beta dispersion) also did not change from Day 1 to Day 8 (ANOVA: F = 1.147, *p* = 0.29).

**Figure 4.**
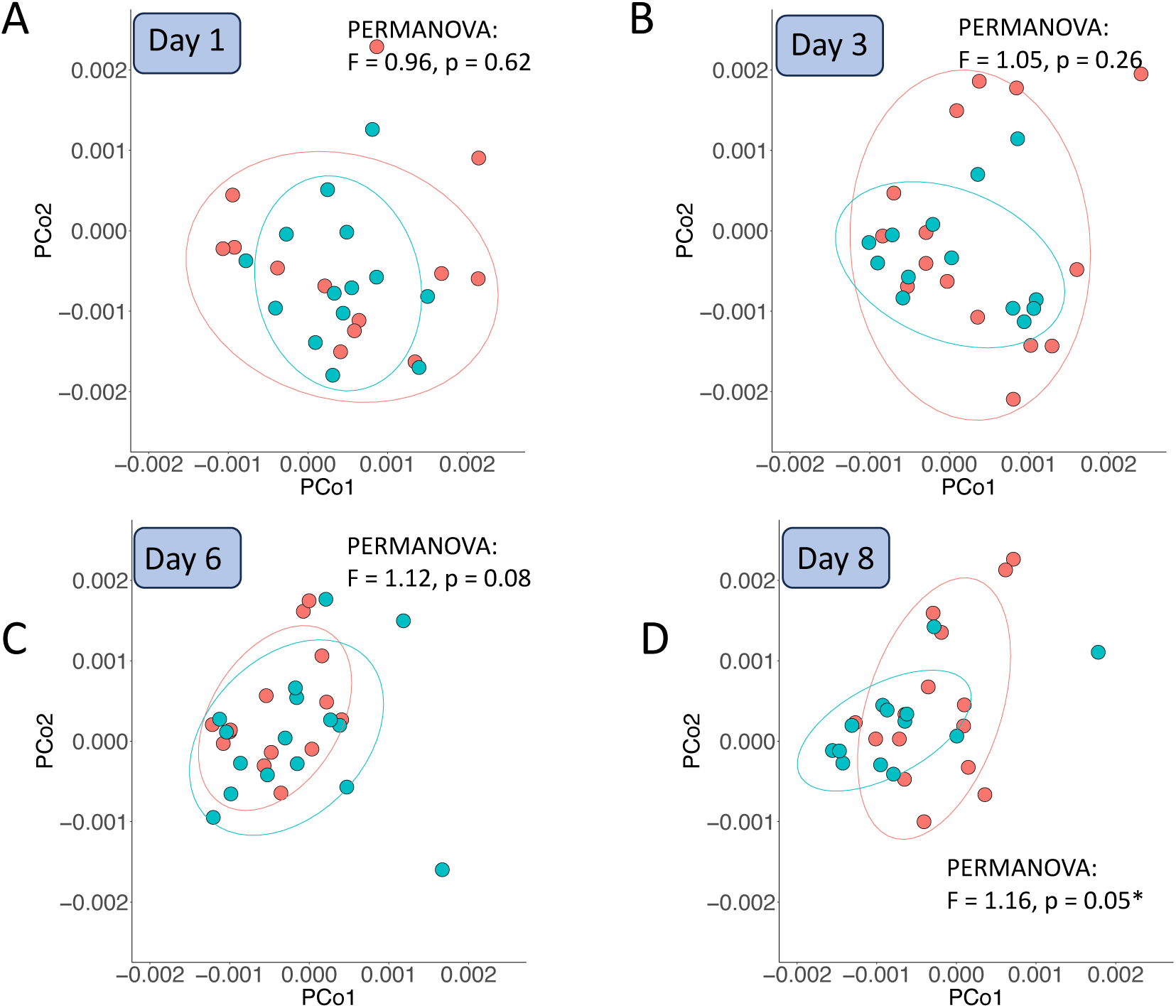
Display of time-series effects on post-exposure community composition between group A and group B, using principal coordinates analysis based on centred-log ratio compositional abundance data for days 1 (panel A), 3 (panel B), 6 (panel C), and 8 (panel D). Red points and ellipses are Group A. Blue points and ellipses are Group B. Outliers were removed on Days 1, 3, and 8.

Time had no effect on alpha diversity for group A (repeated measures ANOVA: ges = 0.084, *p* = 0.30) or group B (repeated measures ANOVA: ges = 0.192, *p* = 0.16; **Figure 5A**). Group B showed a time effect on Gammaproteobacteria diversity, with significantly reduced diversity from Day 1 post to Day 8 post (repeated measures ANOVA: ges = 0.344, *p* = 0.002; **Figure 5B**), but showed no effect on Group A (repeated measures ANOVA: ges = 0.082, *p* = 0.31).

**Figure 5.**
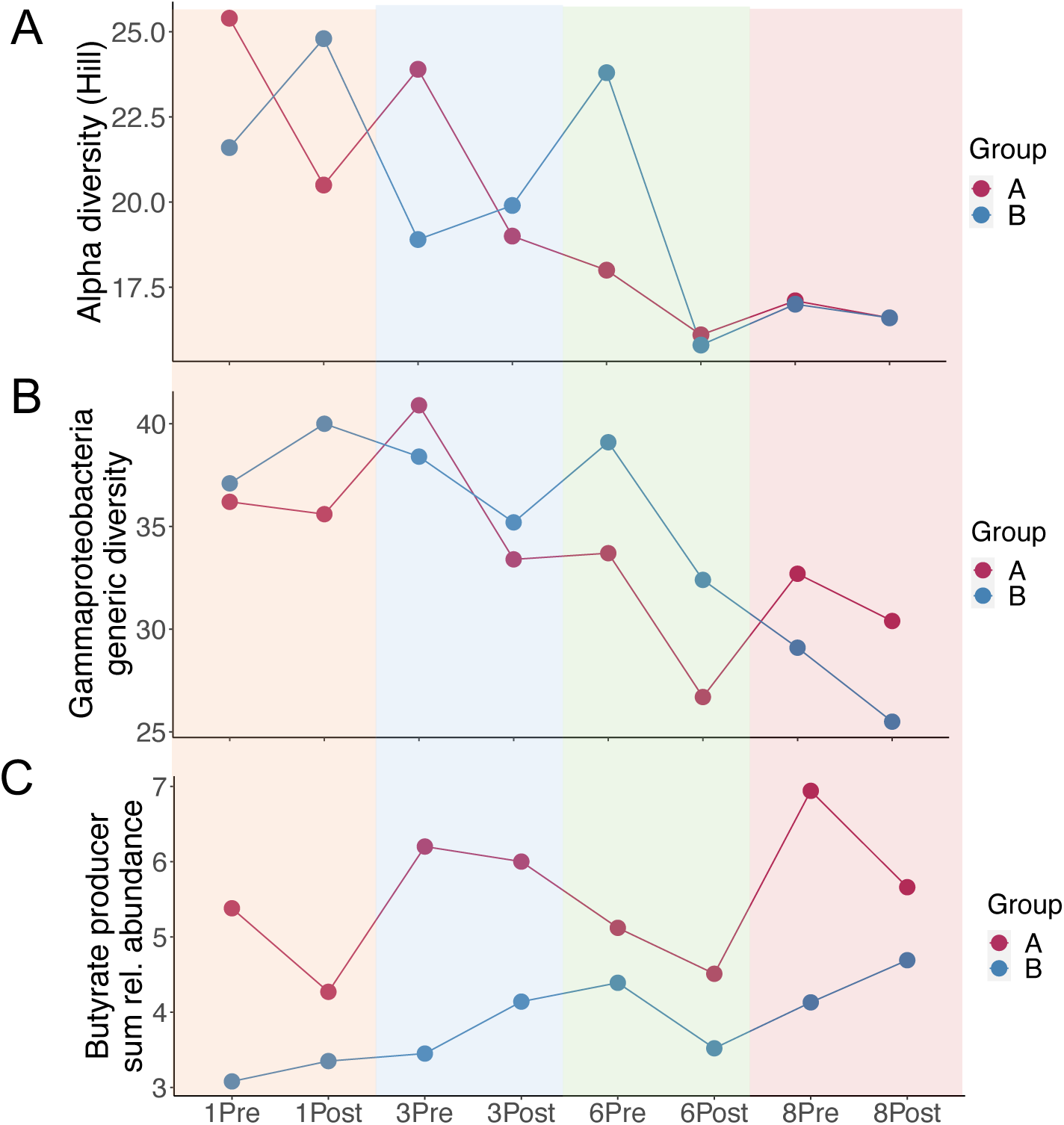
Line plots showing pre-versus post-exposure measures of human health-associated bacterial characteristics in nasal samples of groups A and B across the trial period: (A) alpha diversity (Hill numbers), (B) Gammaproteobacterial generic diversity, and (C) sums of relative abundances of butyrate-producing bacteria.

Time had no effect on the sum of relative abundances of butyrate-producing bacteria for group A (repeated measures ANOVA: ges = 0.119, *p* = 0.14) or B (repeated measures ANOVA: ges = 0.2, *p* = 0.24), although time had a weak effect on increasing read abundances of butyrate-producing bacteria from Day 1 post to Day 8 post in group B (repeated measures ANOVA: ges = 0.167, *p* = 0.058); **Figure 5C**).

## DISCUSSION

We ran a short-term greenspace cross-over exposure trial of a Māori cohort and showed that this exposure had little effect on nasal microbiomes. This low responsiveness of the nasal microbiome was following repeated 30-minute passive exposures to an outdoor nature park. Location, participant, and time had weak or no effect on the nasal microbiome alpha diversity, community composition, aerobiome taxa present in nasal samples, and health-associated bacterial groups. Overall, our results contrast with an earlier study that reported changes in nasal microbiomes after greenspace exposure (11). We suggest that nasal microbiomes are relatively stable over short periods of passive greenspace exposure, and 30 minutes of this passive exposure (i.e., walking in greenspaces) does not result in notable and/or consistent changes in the nasal bacterial communities of participants. Our work raises important questions about the types of activities and duration of exposure to greenspaces required to result in meaningful changes to the nasal microbiome.

### Aerobiomes had higher alpha diversity than nasal microbiomes

We found that overall aerobiomes had higher alpha diversity than nasal microbiomes. This is consistent with the findings from Selway et al. (11), where outdoor air samples had higher alpha diversities than nasal samples. To our knowledge, no previous studies have compared the aerobiome alpha diversity of indoor office and urban greenspace environments. Our findings showed no difference between office and amenity park aerobiome alpha diversity; however, we had only seven air samples (three outdoor and four indoor), which likely limited our power to detect an effect. Recent studies have placed value on urban greenspaces and natural outdoor locations as environmental reservoirs of immunoregulatory biodiversity for urban residents (10, 20). However, future direct comparisons of indoor and outdoor aerobiomes across a range of built environments and outdoor settings are needed to establish the conditions that may drive potential health-promoting exposure effects.

### Short greenspace exposures had little effect on nasal microbiomes

We found no clear effects of the 30-minute exposures on nasal microbiome alpha diversity or community composition. Even when filtering the microbial taxa to just particular health-associated bacterial groups (i.e., Gammaproteobacteria, butyrate-producing bacteria), the only notable effects were a reduction in generic diversity of Gammaproteobacteria and an increase in butyrate-producing bacterial read abundances in group B across the trial period. Roslund et al. (20) recently found that generic diversity of Gammaproteobacteria on the skin of children associated with shifts in blood plasma markers TGF-β1 and interleukin-17 toward an anti-inflammatory profile. Our observed reduction in generic diversity of Gammaproteobacteria and an increase in butyrate-producing bacterial read abundances may be part of normal temporal bacterial variability (37) or could have been driven by an unmeasured factor. However, since so few studies have generated data directly comparable to ours, the capacity to compare our findings with other studies is limited.

### Exposure times

Our trial ran for eight days, with four 30-minute exposure events across these days. We found only minimal changes in nasal microbiome characteristics after each exposure. Our 30-minute exposure length was intended to represent a typical nature exposure of, for example, going for a walk in a park during a lunch break or walking a pet. Similar human exposure trials are limited, but some provide noteworthy discussion. In a study with two or three participants spending time in urban greenspaces, Selway et al. (11) found skin and nasal microbiome changes, but participants performed activities that encouraged more direct interaction with soils and/or vegetation and utilised ca. 1.5 hour exposure periods. Roslund et al. (20) added biodiverse forest floor and sod into daycare centres, then found changes in the skin and gut microbiomes of participant children (3-5 years old) over 28 days with approximately 1.5 hour daily exposure periods. Lai et al. (38) examined the exposure impacts of academic mouse researchers working in the dirty cage wash area on nasal and skin microbiomes. Their exposure period was a single 8-hour shift, and they found no significant change in the nasal microbiome between pre- and post-shift samples. Studies assessing the effects of land cover surrounding a person’s home on their skin microbiome are able to integrate much longer exposure periods to show effects on residents’ microbiomes. For example Hanski et al. (39) assessed the influence of living near biodiversity and found notable effects on the bacterial classes in the skin. Thus, longer and/or repeated exposure periods plus more direct exposure (e.g., handling soils) may be required to elicit changes in nasal and skin microbiomes. Future urban greenspace research should further examine the effect of different activities (e.g., passive walking as in our study, direct handling of microbially-rich ecosystem components such as soil), durations (e.g., short 30-minute periods as in our study, longer and/or repeated short exposures) as well as adjacency and ecological quality of greenspaces on causing changes to human nasal microbiomes.

### Individual participant nasal microbiome stability

We showed a relatively stable participant nasal microbiome over our study period, with strong between-subject effects found on all days. This finding corroborates with human gut microbiome studies, that are generally stable over time (40). Costello et al. (41) described how the host shapes the microbiota through environmental selection processes. We show that the composition of an individual’s nasal microbiome appeared to change over the eight-day period, but not in ways that could be explained by our environmental exposure treatments. Our groups rotated through the same two sites, with similar exposures to the associated aerobiomes. The stability of between-subject microbiome diversity provides additional evidence that more direct, longer, and/or more frequent exposure is necessary for environmental exposures to overcome other host selection pressures to modulate an individual’s nasal microbiome.

### Conclusions

Spending time in urban greenspaces can provide a person with exposure to outdoor aerobiomes that may have health-beneficial properties, such as by providing exposure to high bacterial diversity (10, 20). Our study utilised pre- and post-exposure bacterial data to identify changes in the nasal microbiome following 30-minute walks in an outdoor urban park. We observed stability of the alpha diversity, community composition, and abundances of specific health-associated bacterial groups across exposure periods and across the trial period. Between-subject differences in nasal microbiomes were maintained during the trial period, although some evidence indicated a reduction in the diversity of Gammaproteobacteria and an increase in butyrate producing taxa. Our results suggest that 30 minutes of passive exposure to greenspaces provides insufficient aerobiome exposure to results in changes in nasal bacterial diversity and communities. Indigenous initiatives, which are driven by Indigenous knowledge and emphasise cultural connection as a motivator, could benefit from the expanding collection of microbiome data to better understand the complex (and holistic) relationship between health and the environment. Our study demonstrates the need for future human exposure trials investigating urban greenspace health benefits to examine the types of activity and duration of exposure.

## Acknowledgements

We would like to acknowledge the contributions by Te Arawa Whānau Ora for their participation, enthusiasm, and on-site leadership. Special acknowledgement goes to the laboratory team members at Auckland University of Technology who lent their time, resources, and expertise during the field work and laboratory portions of the project. We would also like to acknowledge Christian Cando-Dumancela for his expertise and assistance in preparing field work resources. This work was supported by funding from the Flinders Foundation and a Project Grant from the Health Research Council of New Zealand.

## Data uploads

All study data and custom R code is available at Figshare at the following doi: 10.6084/m9.figshare.24993471

(Note to Reviewers: We have reserved the DOI mentioned above which will be published upon acceptance; however, for review purposes please see this private figshare link: https://figshare.com/s/78d56add67e2a92db9fb)

## Author contributions

J.B., I.W., D.H., C.L., and M.B. designed and planned the project. J.B., C.L., and M.B. analysed the data, with input from I.W. and D.H. J.B. wrote the initial manuscript, with subsequent revision input from all authors.

