## Supplementary Information for "Short-term passive greenspace exposures have little effect on nasal microbiomes: a cross-over exposure study of a Māori cohort"

**Table S1**. Differential abundances of genera in groups A and B following outdoor exposures. See Figshare (doi: 10.6084/m9.figshare.24993471) for the Excel workbook, with data from each exposure day on a different sheet.

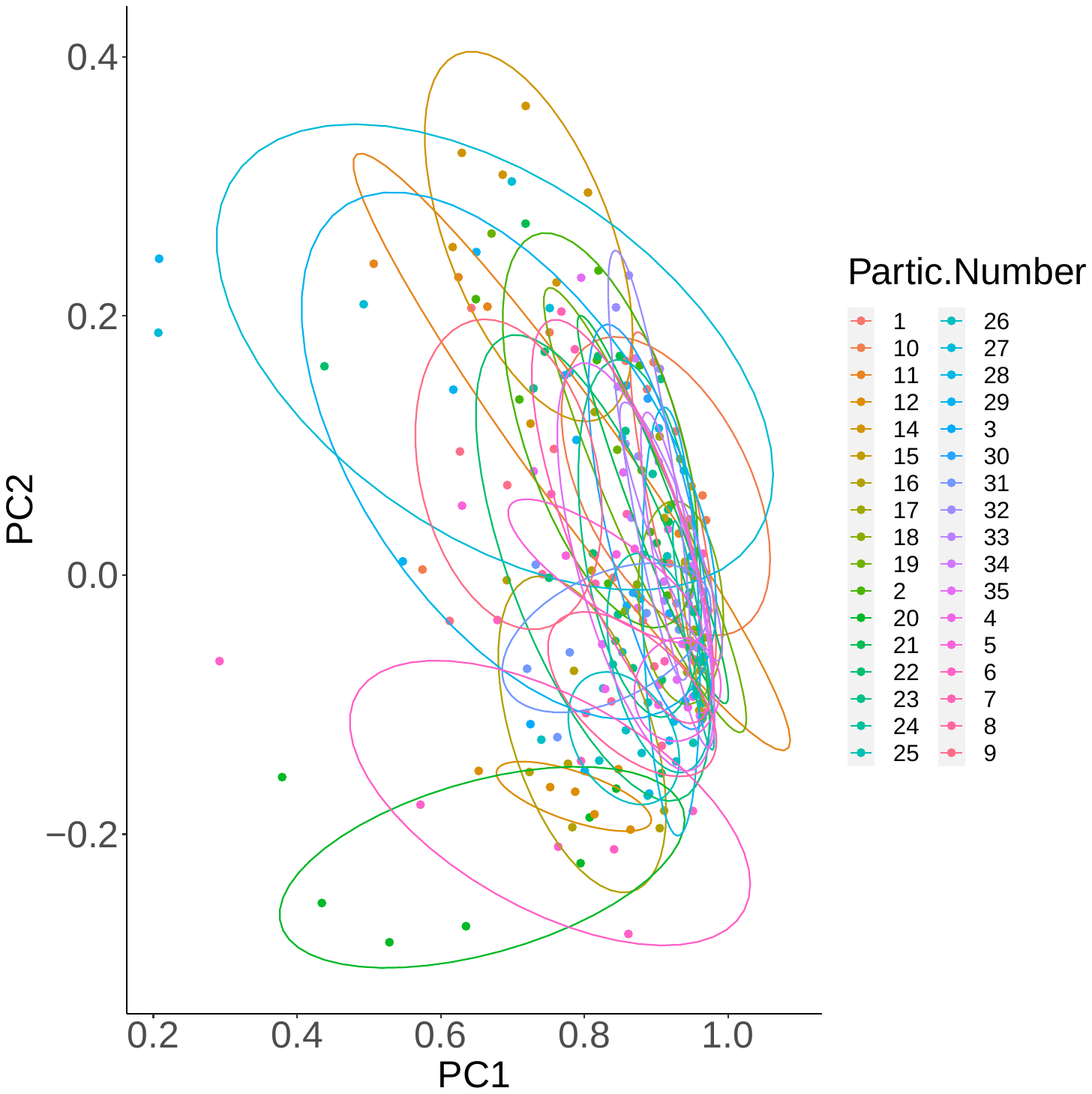

**Figure S1**. Principal components analysis based on centred-log ratio compositional abundance data showing strong effect of participant on the community composition of samples from Day 1 to Day 8 (Adonis PERMANOVA: F = 3.667, R^2^ = 0.382, *p* = 0.001).
